# The MSP-RON axis stimulates cancer cell growth in models of triple negative breast cancer

**DOI:** 10.1101/2020.02.19.956508

**Authors:** Rhona Millar, Anna Kilbey, Sarah-Jane Remak, Tesa M. Severson, Sandeep Dhayade, Emma Sandilands, Kyla Foster, David M. Bryant, Karen Blyth, Seth B. Coffelt

**Affiliations:** Institute of Cancer Sciences, University of Glasgow, Glasgow, United Kingdom; Cancer Research UK Beatson Institute, Glasgow, United Kingdom; Division of Oncogenesis, Netherlands Cancer Institute, 1066CX Amsterdam, The Netherlands

## Abstract

Triple negative breast cancer is the most aggressive subtype of breast cancer with poor prognosis and high rates of relapse. The lack of actionable targets for TNBC has contributed to the high mortality rates of this disease, and new candidate molecules for potential manipulation are urgently required. Here, we show that macrophage-stimulating protein (MSP) and its tyrosine kinase receptor, RON, are potent drivers of cancer cell growth and tumor progression in a mouse model of TNBC driven by the loss of *Trp53* and *Brca1*. After comparison of two genetically engineered mouse models of TNBC, we found that mammary tumors from *K14-Cre;Brca1*^F/F^*;Trp53*^F/F^ (KB1P) mice exhibit high endogenous levels of MSP and RON expression. We show that MSP stimulates AKT and ERK1/2 activation as well as cancer cell growth in KB1P cell lines, while genetic and pharmacological inhibition of RON prevents these effects. Similarly, KB1P tumor progression in mice was robustly attenuated by treatment with a RON inhibitor with accompanied reduction in the proliferation marker, Ki-67. Our findings in a mouse model where MSP and RON expression are naturally increased provide evidence that this receptor and its ligand are viable candidate molecules for targeted treatment of TNBC.

## 1. Introduction

Triple negative breast cancer (TNBC) represents a subgroup of about 15-20% of breast tumors that lacks expression of three main biomarkers: estrogen receptor (ER), progesterone receptor and epidermal growth factor receptor 2 (HER2) amplification (1). When compared to other breast cancer subgroups, patients with TNBC generally have an unfavorable prognosis due to the aggressive nature of this disease and higher risk of local and distant recurrence (2). At the molecular level, TNBC is highly heterogeneous, but unlike ER-positive and HER2-amplified breast cancers, the discovery of uniform actionable molecular targets has eluded researchers (1). Thus, the absence of approved targeted therapies for TNBC leaves chemotherapy as the only therapeutic option.

Recepteur d’origine nantais or RON is a receptor tyrosine kinase encoded by the *Mst1r* gene that shares high homology with the cMET oncogene (3). RON is translated as a single-chain cytoplasmic pro-protein and then further processed to a mature form, which is displayed as a transmembrane protein at the cell surface. Additionally, several isoforms of the RON protein have been reported that stem from two distinct promoters upstream or within the *Mst1r* gene (3). These isoforms can be constitutively active, oncogenic or biologically inactive (3). RON has only one known ligand called macrophage-stimulating protein (MSP). MSP is encoded by the *Mst1* gene, and it is structurally similar to the cMET ligand, hepatocyte growth factor (HGF). RON is lowly expressed in normal mammary gland tissue, but can be overexpressed and phosphorylated in primary breast tumors (4). MSP and RON (over)-expression in breast cancer specimens correlates with poor prognosis and distant recurrence (5, 6), suggesting that the MSP-RON axis contributes to disease progression. Indeed, experimental models, in which MSP or RON are overexpressed or the RON receptor is inhibited or genetically deleted, show that the MSP-RON axis drives mammary tumor progression and metastasis (6–14).

Here, we provide the first evidence of the importance of the MSP-RON axis in a clinically relevant, autochthonous mouse model of TNBC. We show that MSP and RON expression is up-regulated in mammary tumors from *K14-Cre;Brca1*^F/F^;*Trp53*^F/F^ (KB1P) mice when compared with tumors from *K14-Cre;Trp53*^F/F^ (KP) mice or normal mammary glands. Novel cell lines generated from KB1P tumors display increased AKT and ERK1/2 activation in response to MSP-mediated RON signaling. Moreover, MSP stimulates proliferation of KB1P cells whereas inhibition of RON signaling decreases cell growth *in vitro* and tumor growth *in vivo*. These data support the notion that the RON receptor is an actionable target in TNBC and provide a new platform to study MSP-RON biology in a physiologically relevant model.

## 2. Materials & Methods

### 2.1 Animal Models and Cell Lines

The generation and characterization of *K14-Cre;Trp53*^F/F^ (KP) and *K14-Cre;Brca1*^F/F^;*Trp53*^F/F^ (KB1P) mice, which are maintained on the FVB/n background, is described here (15). Single cell suspensions from end-stage (15mm) mammary tumors were plated and propagated in hypoxic incubators for four weeks to generate cell lines. These cells were grown in DMEM supplemented with 10% fetal calf serum (FCS), 100U/mL penicillin/streptomycin and 2mM glutamine. Before recombinant MSP treatment (100ng/mL; R&D systems #6244-MS-025), cells were serum starved in 0.2% FSC overnight. The RON inhibitor BMS-777607 (ChemieTek #CT-BMS777) was added to cells at 1μM for 1h before MSP treatment where indicated.

The orthotopic transplantation of tumor fragments is described here (16). Briefly, KB1P tumor fragments were implanted into the fourth mammary fat pad of 10-12 week old female recipient FVB/n mice (Charles River, UK). When tumors reached 3×3 mm, mice were given a daily oral gavage of BMS-777607 (50mg/kg dissolved in DMSO and diluted in 70% PEG-300) or vehicle control (11% DMSO diluted in 70% PEG-300) and then monitored until tumors reached 15 mm.

Mouse experiments were performed in accordance UK Home Office license numbers 70/8645 (Karen Blyth), carried out in-line with the Animals (Scientific Procedures) Act 1986 and the EU Directive 2010, and sanctioned by Local Ethical Review Process (University of Glasgow). Mice were housed on a 12/12 light/dark cycle and fed and watered ad libitum.

### 2.2 RNA sequencing analysis

Total tumor gene expression data from *K14-Cre;Trp53*^F/F^ (KP) and *K14-Cre;Brca1*^F/F^;*Trp53*^F/F^ (KB1P) mice were kindly provided by Lewis Cantley (NCBI BioProject #PRJNA398328). RNA isolation, DNA library preparation, sequencing information and generation of FPKM values for these data is described here (17). To visualize the gene expression FPKM values were log transformed in R version 3.5.0 and boxplots of generated with the ggplot2 package. Differences between expression distributions were tested with the Wilcoxon rank sum test.

### 2.3 Quantitative real-time PCR

RNA was isolated from frozen mammary tumors or cell lines using Qiagen’s RNeasy kit and on-column DNA digestion. RNA concentration was determined using a Thermoscietific NanoDrop spectrophotometer with NanoDrop 2000 software. cDNA was prepared from 1 μg RNA using a Quantitect Reverse Transcription kit (Qiagen) and diluted 1:20 in DEPC-treated water. For quantitative real-time PCR, 12.5 ng aliquots of cDNA were amplified in triplicate on an ABI 7500 real-time PCR machine using SyGreen Blue Mix Lo-ROX PCR master mix (PCRBIOSYSTEMS) and primers for *Mst1r (*Mm_Mst1r_1-SG; Quantitect) and *Mst1 (Forward – CTCACCACTGAATGACTTCCAG; Reverse - AAGGCCCGACAGTCCAGAA)* at 2.5μM with endogenous controls *Hprt*(Mm_Hprt_1_SG; Quantitect) and *β*-actin (Mm_Actn_1_SG; Quantitect). Relative expression was calculated by the ΔCt method and data are displayed as 1/ΔCt.

### 2.4 Western blotting

Protein was extracted from mammary tumors or cell lines using RIPA buffer (50 mM Tris-HCl, pH 7.4, 150 mM NaCl, 1% NP-40, 0.5% DOC, 0.1% SDS, 2 mM EDTA) complemented with 1x protease/phosphatase inhibitor cocktail (HALT 100x, Thermo Fisher Scientific). Mammary tumor tissue was physically disrupted in the same mix with Precellys Hard Tissue beads (VWR) using a rotor-stator homogenizer until uniformly homogenous. Lysates were clarified by centrifugation and protein levels were quantified using a BCA protein assay kit (Thermo Fisher Scientific). Proteins (50 μg) were resolved on 4-12% NuPage polyacrylamide gels (Invitrogen) and transferred onto enhanced chemiluminescence (ECL) nitrocellulose membranes using an iBlot2 Transblot Turbo Transfer System (Invitrogen). Primary antibodies were incubated overnight at 4^0^C on blocked membranes: anti-RON (500ng/mL, Thermo Fisher Scientific, #PA5-71878,); anti-MSP (1μg/mL, R&D Systems, #AF6244); lnti-phospho-ERK1/2 (250ng/mL, Cell Signaling Technology, #4370); anti-ERK1/2 (84ng/mL; Cell Signaling Technology, #4695); anti-AKT (34ng/mL, Cell Signaling Technology, #9272); anti-phospho-AKT (17ng/mL, Cell Signaling Technology, #4058); and 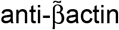 (200ng/mL, Sigma-Aldrich, #A5316). HRP linked secondary antibodies (Cell Signaling Technology) were incubated for 1h at room temperature and proteins visualized by chemiluminescence (Thermo Fisher Scientific). Each experiment was repeated at least four times.

### 2.5 ELISA

MSP serum levels from autochthonous, mammary tumor-bearing KP and KB1P mice or tumor-free, wild-type mice were measured using a kit from R&D Systems according to the manufacturer’s recommendations.

### 2.6 Lentiviral transduction

Target sequences specific for *Mst1r* were designed using the iRNAi program. shRNA oligos were generated and sub-cloned into the pLKO.1puro lentiviral backbone using Addgene’s protocol (https://www.addgene.org/8453/). Viral supernatants were prepared following transient transfection of 293FT cells with pLKO.1 encoding shRNAs, pSPAX2 packaging vector and pVSVG envelope vector using Lipofectamine 2000 according to the manufacturer’s instructions. Two 24h supernatants were collected sequentially over a 48h period, pooled and filtered through a 0.45μm syringe filter and then concentrated using the Lenti-X Concentrator solution (Clontech). Freshly concentrated supernatants were added directly to drained sub-confluent recipient cells and incubated overnight. An equal volume of fresh medium was then added and antibiotic selection (puromycin) initiated 48h later. Antibiotic resistant clones were expanded for further analysis. We tested 9 different shRNA oligos. Efficient *Mst1r* knockdown was determined by quantitative real-time PCR and two distinct shRNAs were selected for further experiments. Target sequences of the two oligos used for this study were as follows: shRON-1 sense - CCTGCTGTATGTGTCCAACTT, anti-sense - AAGTTGGACACATACAGCAGG; shRON-2 sense - CGTCCTAGACAAGGAATACTA, anti-sense – TAGTATTCCTTGTCTAGGACG.

### 2.7 Proliferation Assay

Cellular proliferation was measured using the fluorescence-based proliferation CyQuant NF kit (Thermo Fisher Scientific) according to the manufacturer’s instructions. Cells were seeded in a 96-well plate, 6 replicates per condition at 10^4^ cells/well in low serum (0.2%) for 72h with daily administration of 100ng/mL MSP and/or 1μM BMS-777607.

### 2.8 Immunohistochemistry

Immunohistochemical analyses were performed by the Histology facility at the CRUK Beatson Institute using standard protocols on Bond Rx or Dako autostainers. Anti-Ki67 (clone SP6; 1:100) was purchased from Abcam and anti-Caspase 3 (clone Asp-175; 1:500) was purchased from Cell Signaling. Quantitative analysis of positive staining was performed by counting cells in at least 3 high-power fields of view (×40) per tumor by two independent researchers who were blinded to the sample group. Images were captured on an Axio Imager A2 Bio upright microscope (Zeiss) using ZenPro 2012 software (Zeiss).

### 2.9 Statistics

The non-parametric Mann-Whitney U test was used to compare two groups, while one-way ANOVA was used to compare groups of three or more. Two-way ANOVA with repeated measures was used to analyze tumor growth curves. The log-rank (Mantel-Cox) test was used to analyze Kaplan-Meier survival curves. Sample sizes for each experiment were based on a power calculation and/or previous experience of the mouse models. Analyses were performed using GraphPad Prism (version 8).

## 3. Results

### 3.1 MSP and its receptor, RON, are expressed in autochthonous TNBC models

In previous studies, gene expression analysis of mammary tumors from *K14-Cre;Trp53*^F/F^ (KP) and *K14-Cre;Brca1*^F/F^;*Trp53*^F/F^ (KB1P) mice by microarray revealed that 646 genes are differentially expressed between these two models of TNBC (15). To identify targetable vulnerabilities in these mammary tumors, we mined this list for indicators of aberrant signaling pathways. We found that expression of the *Mst1* gene – which encodes the RON receptor ligand, macrophage stimulating protein (MSP) – is increased in KB1P tumors when compared with KP tumors. We analyzed RNA sequencing data from KP and KB1P tumors for expression levels of *Mst1* and *Mst1r* (17). *Mst1* levels were increased in KB1P tumors when compared to normal mammary gland tissue and KP tumors, while *Mst1r* levels were lower in both KP and KB1P tumors when compared to normal mammary gland (Fig. 1A). We validated these findings by real-time PCR. *Mst1* mRNA levels were significantly higher in KB1P tumors, but *Mst1r* levels were the same between tumor models (Fig. 1B). Expression levels of MSP protein mirrored the mRNA levels and were greater in KB1P compared to KP derived tumor tissues (Fig. 1C). Similarly, MSP serum levels were also elevated in mammary tumor-bearing KB1P mice, when compared to tumor-bearing KP mice or tumor-free, wild-type (WT) mice (Fig. 1D). Western blot analysis of RON expression revealed multiple isoforms of the RON receptor (Fig. 1E). The short form of RON was most prevalent and, in keeping with the qRT-PCR analysis, was expressed to approximately equivalent levels in both tumor models. In contrast, the long form (RON β chain) was differentially expressed between KP and KB1P tumors, where 4 out of 5 KB1P tumors expressed full length RON-β, but only 1 out of 5 KP tumors showed low level expression of RON-β Of the remaining 4 KP tumors, 3 tumors expressed pro-RON, the inactive form of the receptor, and 1 tumor expressed no detectable long-forms of RON (Fig. 1e). Quantification of the combined full length RON band intensities revealed overall higher expression levels in KB1P compared to KP tumors (Fig. 1E). Together these data reveal that the MSP specific form of the RON receptor was differentially expressed between KP and KB1P tumor models, and the short form of RON, which lacks the MSP binding domain and remains unresponsive to MSP, was present at high levels in both tumor models.

**Figure 1.**
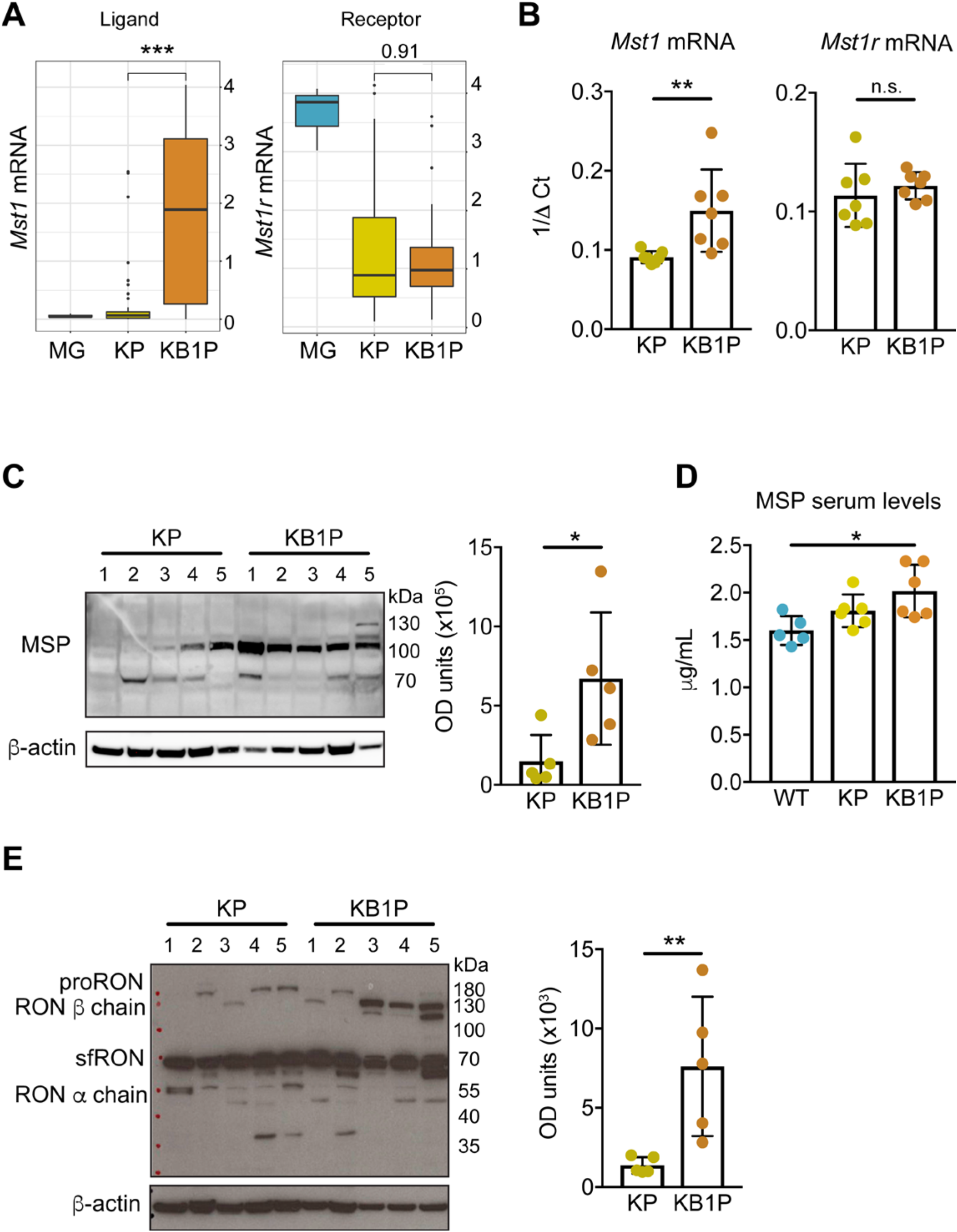
MSP and RON are upregulated in tumors from KB1P mice. (A) *Mst1* and *Mst1r* mRNA expression in normal mammary glands (MG) (from 3 individual mice), KP tumors (n=44) and KB1P tumors (n=41) analyzed by RNA sequencing. (B) *Mst1* and *Mst1r* mRNA expression in KP and KB1P tumors assessed by qRT-PCR. (C) Western blot analysis of MSP protein expression in KP and KB1P tumors with densitometric quantification. Numbers above the blot represent individual tumors. (D) MSP serum levels in tumor-free, wild-type (WT) mice, tumor-bearing KP mice and tumor-bearing KB1P mice assessed by ELISA. (E) Western blot analysis of RON protein expression in KP and KB1P tumors with densitometric quantification. Numbers above the blot represent individual tumors. Each dot represents one donor tumor from one mouse. \**p*<0.05, \*\**p*<0.01, \*\*\**p*<0.001 as determined by Mann-Whitney U test or one-way ANOVA.

### 3.2 MSP-RON signaling activate AKT and MAPK pathways

We derived cell lines from four independent KP and KB1P tumors, and performed qRT-PCR analysis to confirm continued expression of *Mst1r* mRNA *in vitro*. As shown in Fig. 2A, *Mst1r* transcripts persisted in all cell lines. Protein expression was then measured by western blot analysis in the same cell lines. In contrast to the primary tumor tissue (Fig. 1E), a single band corresponding to the full length RON receptor β chain was the predominant species detected and it was expressed to approximately equivalent levels in each cell line (Fig. 2B). Expression of the short form of RON also persisted, but the ratio between the long and short forms had increased compared to the primary tumors. Given that MSP and full length RON protein were increased in tumors from KB1P mice (Fig. 1), we focused solely on this model for functional experiments and signaling pathways downstream of MSP/RON. To this end, two independent KB1P cell lines were treated with recombinant MSP and protein lysates were analyzed for activation of AKT and ERK1/2. As shown in Fig. 2C, MSP treatment induced robust signaling responses, inducing phosphorylation of AKT and ERK1/2 in both cell lines. To confirm that MSP is acting through the RON receptor, KB1P cells were pre-incubated with a RON inhibitor, BMS-777607. For both cell lines, pre-incubation with BMS-777607 effectively reduced phosphorylation of AKT and ERK1/2 in response to MSP stimulation (Fig. 2C). To confirm selectivity against RON (since BMS-777607 is not absolutely specific for the RON receptor), we repeated these experiments using an shRNA-mediated knockdown approach to silence *Mst1r* expression (Fig. 2D). Mirroring pharmacological inhibition of RON, MSP treatment of KB1P cells lacking RON failed to increase phosphorylation of both AKT and ERK1/2 (Fig. 2E). These data indicate that MSP stimulates robust intracellular signaling pathways via the RON receptor on KB1P cells.

**Figure 2.**
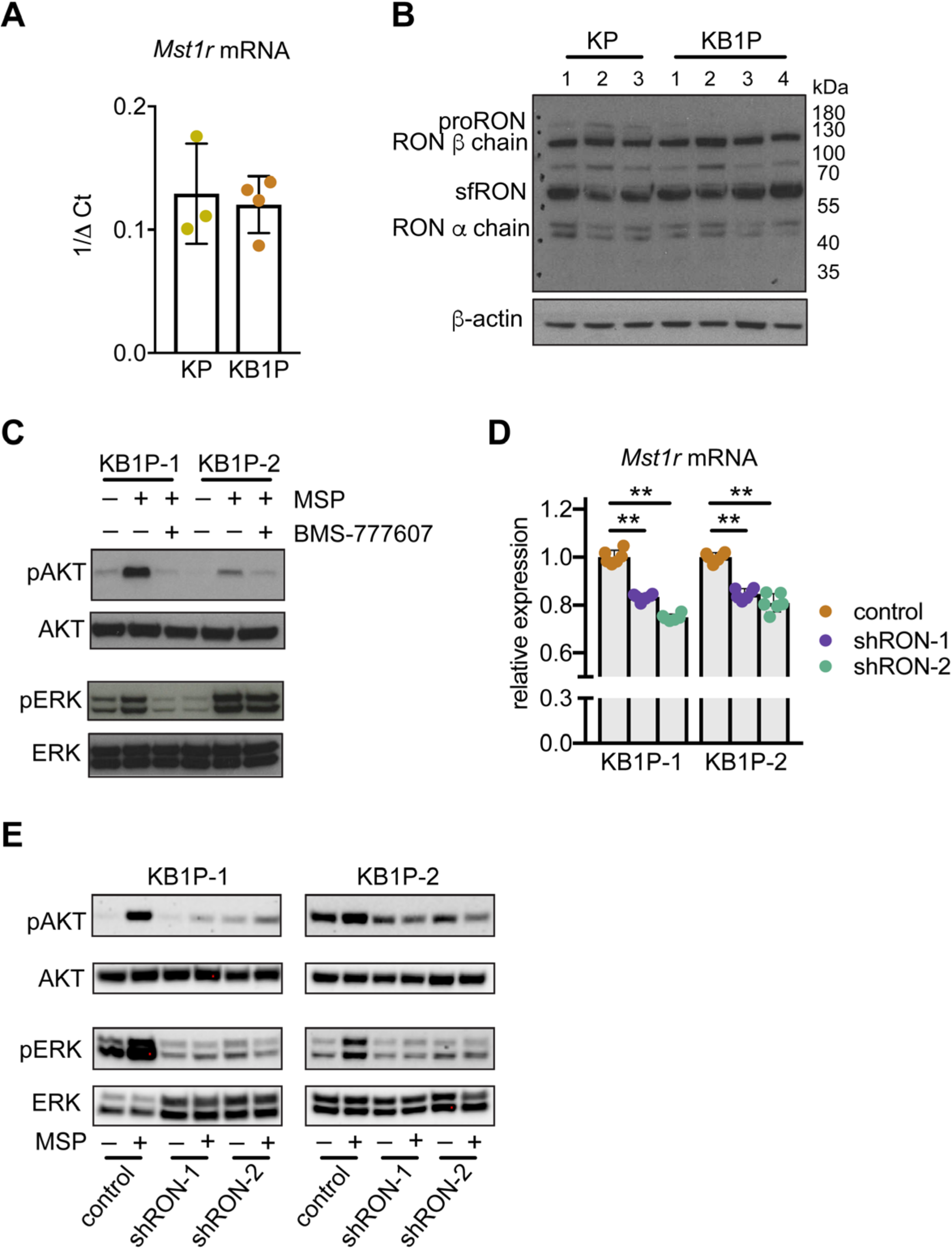
MSP stimulates AKT and MAPK signaling pathways through the RON receptor. (A) *Mst1r* mRNA expression assessed by qRT-PCR in cell lines derived from three KP tumors and four KB1P tumors, respectively. (B) Western blot analysis of RON protein expression in the same cell lines used in A. (C) Western blot analysis of indicated proteins in two KB1P cell lines. Cells were pre-treated with 1μM BMS-777607 for 1h prior to 100ng/mL recombinant MSP for 1h where denoted. Images are representative of three replicates. (D) Two KB1P cell lines were transduced with lentiviral shRNA vectors against *Mst1r* mRNA (shRON-1 or shRON-2) or control pLKO.1 vector. Confirmation of *Mst1r* mRNA knockdown quantified by qRT-PCR is expressed as relative to two housekeeping genes, *Hprt* and *β*-actin (each dot represents one technical replicate in a given experiment, the experiment was repeated three times for each cell line). (E) Two independent KB1P cell lines transduced with control or shRNA vectors against *Mst1r* mRNA were treated with 100ng/mL recombinant MSP for 1h. Activation of AKT and ERK1/2 was assessed by western blot. Images are representative of three replicates. \*\**p*<0.01 as determined by one-way ANOVA.

### 3.3 The MSP-RON axis promotes cell proliferation

We examined the functional effects of MSP-RON signaling on proliferation of two independent KB1P cell lines, since both pathways are known to regulate cell division. We found that MSP increased proliferation of both cell lines, and that proliferation could be attenuated by addition of BMS-777607 (Fig. 3A). Treatment of either KB1P cell line with BMS-777607 alone had no effect on cell proliferation compared with untreated control cells (Fig. 3A). Moreover, shRNA-mediated RON depletion, using two shRNAs separately in each KB1P cell line, significantly reduced proliferation after MSP treatment (Fig. 3B). These data indicate that MSP requires long form RON for its mitogenic effects.

**Figure 3.**
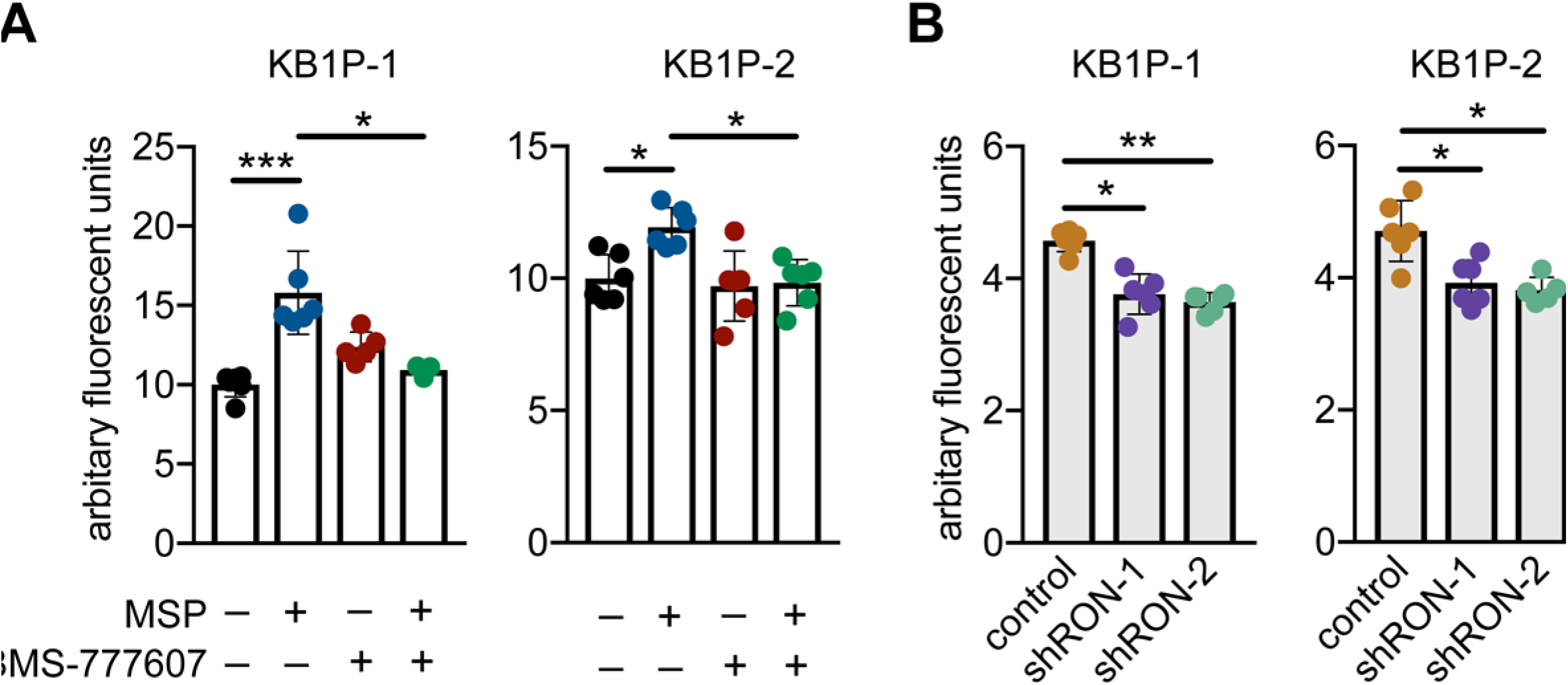
MSP activation of the RON receptor induces proliferation of KB1P cells *in vitro* and tumor growth *in vivo*. (A) Proliferation of two KB1P cell lines in response to 100ng/ml MSP, 1uM BMS-777607 or both reagents. For combined treatments cells were pre-treated with BMS-777607 for 1h before the addition of MSP. For all treatments cells were re-fed every 24h with fresh MSP and BMS-777607 as denoted. DNA content was measured at 72h post treatment. (B) Proliferation of two KB1P cell lines transduced with control pLKO.1 or shRNA vectors against *Mst1r* mRNA following 48h treatment with recombinant MSP as above. For both A and B, each dot represents one technical replicate from one experiment. The experiment was repeated three times. \**p*<0.05, \*\**p*<0.01, \*\*\**p*<0.001 as determined by one-way ANOVA.

### 3.4 Inhibition of RON delays tumor growth in vivo

Next, we transplanted tumor fragments from KB1P mice into the mammary glands of syngeneic FVB/n mice. When tumors were palpable, we treated the mice with BMS-777607 or vehicle control and monitored tumor growth until humane end-point. Inhibition of RON in KB1P mammary tumor-bearing mice delayed tumor growth. This difference was statistically significant (*p* < 0.01) beginning from day 13 (Fig. 4A). In addition, KB1P tumor-bearing mice treated with BMS-777607 survived significantly longer (7 days) when compared to the control cohort (Fig. 4B). To gain insight into how RON signaling potentiates tumorigenesis, mice transplanted with KB1P tumor fragments were treated mice with BMS-777607 or vehicle control for 7 days and then euthanized to examine the growing tumors histologically. The number of Ki-67+ cells and Caspase 3+ cells were measured as surrogate markers for proliferation and apoptosis, respectively..Tumor sections from BMS-777607-treated mice exhibited less Ki-67+ cells when compared to controls (Fig. 4C), whereas Caspase 3+ cells were similar between both groups (Fig. 4D). Together these data indicate that RON inhibition slows cancer cell proliferation without affecting apoptosis and suggest that, in the KB1P model, the MSP-RON axis plays an important role in cancer progression.

**Figure 4.**
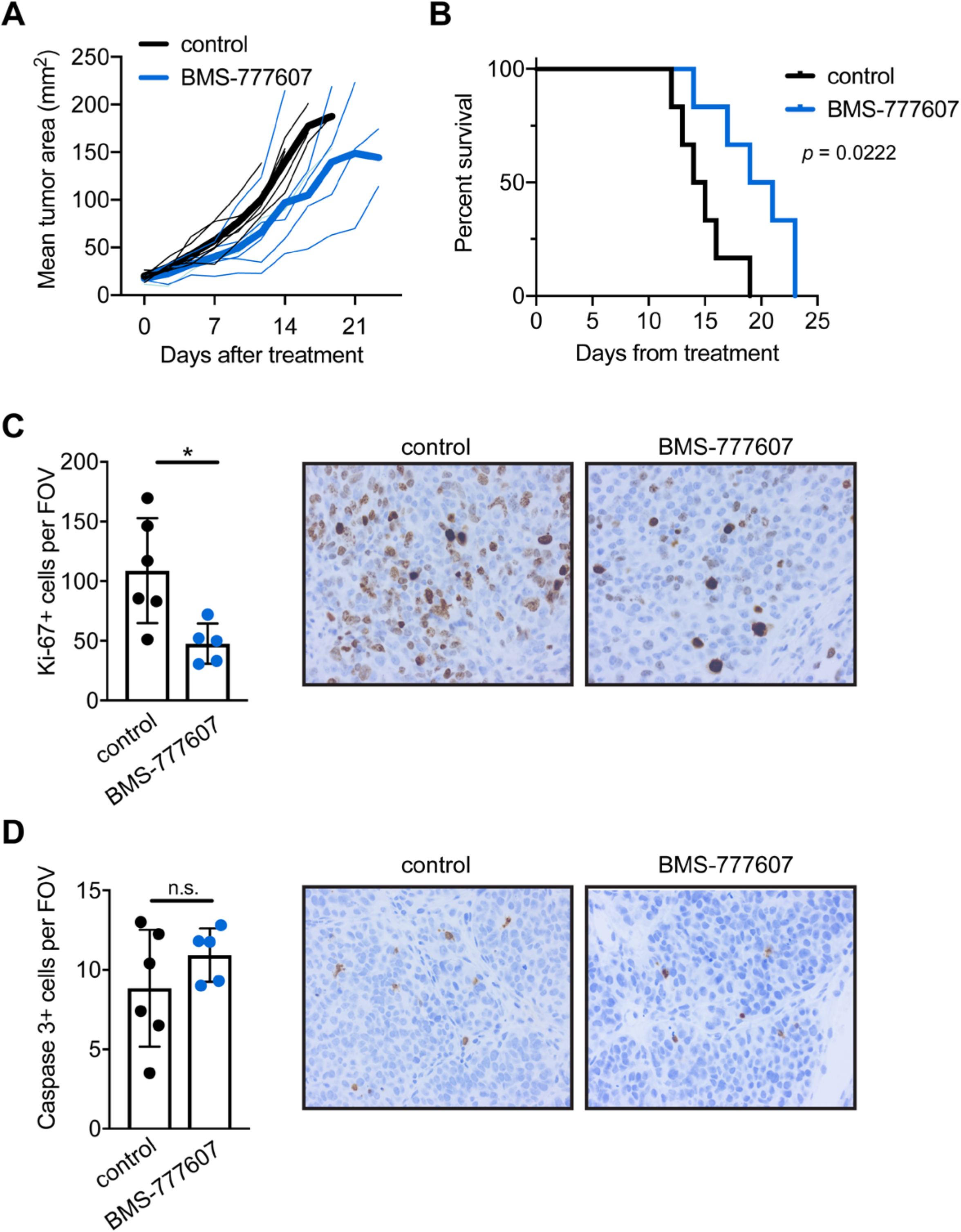
Inhibition of RON signaling reduces KB1P tumor growth in mice. Mice were transplanted with KB1P tumor fragments. Once tumors reached 3×3mm, mice were treated daily with 50mg/kg BMS-777607 or vehicle control for 10 days in A, B (n=6/group) and 7 days in C, D (n=5-6/group). (A) Tumor growth was measured by calipers twice per week and represented graphically. Dark solid lines indicate the mean of each group, while thin lines indicate individual mice. Black=control, blue=BMS-777607. Tumor growth was significantly different (*p* < 0.01) between groups beginning on day 13. (B) Kaplan-Meier survival analysis of the mice depicted in C euthanized when tumors reached 15×15mm in size. Statistical difference was calculated by Mantel-Cox test. (C-D) Tumor sections were processed by immunohistochemistry for (C) Ki-67 and (D) cleaved Caspase 3. Staining was quantified by counting positive cells in at least three 40X fields of view. Each dot represents one tumor. Representative images are shown for each stain from control or BMS-777607-treated tumors. \**p*<0.05 as determined by Mann-Whitney U test.

## 4. Discussion

In this study, we found that MSP signaling via its receptor, RON, is critical for growth and proliferation of KB1P mammary cancer cells both *in vitro* and *in vivo*. These data accord well with previous publications in other mammary tumor models that rely on artificial, ectopic expression of MSP or the RON receptor (6, 7, 9–12, 14). Most mechanistic work on the MSP-RON axis has used cell lines derived from the *MMTV-PyMT* model (6, 9, 12, 14) that over-express MSP, or genetically engineered mouse models in which *Mst1r* expression is driven by the *MMTV* promoter (7, 11). The KB1P model provides a new resource to interrogate the importance of MSP and RON in a non-viral oncogenic model that better mimics breast cancer etiology.

Our findings provide additional insight into the role of RON isoforms in cancer progression. We show that primary mammary tumors from the KP and KB1P models express multiple isoforms of RON with the short form being the most dominant. Full-length RON and short-form RON mRNA are initiated from different transcriptional start sites (3); although, the regulation of these isoforms in KP and KB1P mammary cancer cells is unknown. When cell lines were generated from these two models, expression of the longer forms of RON equaled the expression of the short form of RON. These data indicate a discrepancy between tumors *in situ* and mammary cancer cell lines grown on plastic that should be considered when designing experiments. This observation also suggests that persistence of the long form of RON provides an advantage to cancer cells grown in culture. Both long and short isoforms of RON are inhibited by BMS-777607. Interestingly, treatment of two independent KB1P cell lines with BMS-777607 alone (in the absence of MSP) failed to block proliferation suggesting that short form RON is less important for cell growth of KB1P cells than the long form, which requires MSP for activation.

Another interesting observation from this work was the upregulation of MSP and RON expression in BRCA1-deficient mammary tumors compared to BRCA1 wild-type tumors, which suggests that MSP/RON signaling is evolutionarily selected in highly genomically unstable tumors. A previous publication exhaustively characterized copy number alterations in tumors from the KP and KB1P models and showed that allele frequency of *Mst1* and *Mst1r* genes is normal in both tumor types, without amplification or deletion (17). Since upregulation of MSP and RON in KB1P tumors cannot be explained by increased copy number it would seem that over-active transcription and/or translation are the most likely cause. Given the importance of the MSP-RON axis in BRCA1-deficient mammary tumors, it will be interesting to investigate the synergy of RON inhibitors with PARP inhibitors to mitigate resistance.

## 5. Conclusion

Our results show that the MSP-RON axis is specifically upregulated in a BRCA1-deficient autochthonous mouse model of TNBC. Pharmacological or genetic inhibition of the RON receptor resulted in attenuated AKT and MAPK signaling and reduced cell growth *in vitro*, whereas manipulation of the pathway significantly improved survival and delayed tumor growth *in vivo*. These data identify a clinically relevant model system that will be useful to study the biology and therapeutic targeting of the MSP/RON signaling pathway *in vivo.*.

## 6. Author contributions

Concept, design and supervision: RM, AK, DMB, KB, SBC. Data acquisition, analysis and interpretation: all authors. Writing and review of manuscript: RM, AK, DMB, KB, SBC. Reading and approval of final manuscript: all authors.

## 7. Acknowledgements and funding sources

The authors thank Jos Jonkers for advice and for providing the mouse models. We thank Lewis Cantley (Meyer Cancer Center, Weill Cornell Medical College, New York) for providing RNA sequencing data. The authors thank the core cervices at the Cancer Research UK Beatson Institute (C596/A17196), with particular thanks to the Beatson Histology facility. The William Forrest Charitable Trust PhD Research Fund and the Cancer Research UK Glasgow Center (A25142) funded this work.

## References

1. Bianchini G, Balko JM, Mayer IA, Sanders ME, Gianni L. Triple-negative breast cancer: challenges and opportunities of a heterogeneous disease. Nature reviews Clinical oncology. 2016;13(11):674–90.

2. Malorni L, Shetty PB, De Angelis C, Hilsenbeck S, Rimawi MF, Elledge R, et al. Clinical and biologic features of triple-negative breast cancers in a large cohort of patients with long-term follow-up. Breast cancer research and treatment. 2012;136(3):795–804.

3. Yao HP, Zhou YQ, Zhang R, Wang MH. MSP-RON signalling in cancer: pathogenesis and therapeutic potential. Nat Rev Cancer. 2013;13(7):466–81.

4. Maggiora P, Marchio S, Stella MC, Giai M, Belfiore A, De Bortoli M, et al. Overexpression of the RON gene in human breast carcinoma. Oncogene. 1998;16(22):2927–33.

5. Lee WY, Chen HH, Chow NH, Su WC, Lin PW, Guo HR. Prognostic significance of co-expression of RON and MET receptors in node-negative breast cancer patients. Clin Cancer Res. 2005;11(6):2222–8.

6. Welm AL, Sneddon JB, Taylor C, Nuyten DS, van de Vijver MJ, Hasegawa BH, et al. The macrophage-stimulating protein pathway promotes metastasis in a mouse model for breast cancer and predicts poor prognosis in humans. Proc Natl Acad Sci U S A. 2007;104(18):7570–5.

7. Zinser GM, Leonis MA, Toney K, Pathrose P, Thobe M, Kader SA, et al. Mammary-specific Ron receptor overexpression induces highly metastatic mammary tumors associated with beta-catenin activation. Cancer Res. 2006;66(24):11967–74.

8. Liu X, Zhao L, Derose YS, Lin YC, Bieniasz M, Eyob H, et al. Short-Form Ron Promotes Spontaneous Breast Cancer Metastasis through Interaction with Phosphoinositide 3-Kinase. Genes Cancer. 2011;2(7):753–62.

9. Eyob H, Ekiz HA, Derose YS, Waltz SE, Williams MA, Welm AL. Inhibition of ron kinase blocks conversion of micrometastases to overt metastases by boosting antitumor immunity. Cancer Discov. 2013;3(7):751–60.

10. Cunha S, Lin YC, Goossen EA, DeVette CI, Albertella MR, Thomson S, et al. The RON receptor tyrosine kinase promotes metastasis by triggering MBD4-dependent DNA methylation reprogramming. Cell Rep. 2014;6(1):141–54.

11. Benight NM, Wagh PK, Zinser GM, Peace BE, Stuart WD, Vasiliauskas J, et al. HGFL supports mammary tumorigenesis by enhancing tumor cell intrinsic survival and influencing macrophage and T-cell responses. Oncotarget. 2015;6(19):17445–61.

12. Andrade K, Fornetti J, Zhao L, Miller SC, Randall RL, Anderson N, et al. RON kinase: A target for treatment of cancer-induced bone destruction and osteoporosis. Sci Transl Med. 2017;9(374).

13. Faham N, Zhao L, Welm AL. mTORC1 is a key mediator of RON-dependent breast cancer metastasis with therapeutic potential. NPJ Breast Cancer. 2018;4:36.

14. Ekiz HA, Lai SA, Gundlapalli H, Haroun F, Williams MA, Welm AL. Inhibition of RON kinase potentiates anti-CTLA-4 immunotherapy to shrink breast tumors and prevent metastatic outgrowth. Oncoimmunology. 2018;7(9):e1480286.

15. Liu X, Holstege H, van der Gulden H, Treur-Mulder M, Zevenhoven J, Velds A, et al. Somatic loss of BRCA1 and p53 in mice induces mammary tumors with features of human BRCA1-mutated basal-like breast cancer. Proc Natl Acad Sci U S A. 2007;104(29):12111–6.

16. Doornebal CW, Klarenbeek S, Braumuller TM, Klijn CN, Ciampricotti M, Hau CS, et al. A preclinical mouse model of invasive lobular breast cancer metastasis. Cancer Res. 2013;73(1):353–63.

17. Liu H, Murphy CJ, Karreth FA, Emdal KB, White FM, Elemento O, et al. Identifying and Targeting Sporadic Oncogenic Genetic Aberrations in Mouse Models of Triple-Negative Breast Cancer. Cancer Discov. 2018;8(3):354–69.

